# Rhizobacteria prime the activation of defence and nutritional responses to suppress aphid populations on barley

**DOI:** 10.1101/2024.09.04.611222

**Authors:** Crispus M. Mbaluto, Sharon E. Zytynska

## Abstract

- Interactions between plant and soil microbes are widespread and modulate plant-insect herbivore interactions. Still, it remains unclear how these shapes the overall plant defence responses and the mechanisms involved.
- Here, we performed bioassays with barley (*Hordeum vulgare*) plants to study the underlying molecular pathways induced by two rhizobacteria, *Acidovorax radicis* or *Bacillus subtilis,* against the phloem feeding aphid *Sitobion avenae* over three timepoints.
- Root colonization by *A. radicis* or *B. subtilis* suppressed aphid populations on barley. Analysis of differentially expressed genes and co-expressed gene modules revealed a combination of rhizobacteria and aphid induced plant responses. Aphid feeding triggered distinct plant responses in rhizobacteria-inoculated barley compared to controls, in phytohormone, glutathione, and phenylpropanoid pathways within 24 hours. By day 7, stronger responses were observed in phenylpropanoid and nutrient pathways. By day 21, changes occurred in flavonoid pathways and genes related to tissue damage and repair.
- Our study suggests that rhizobacteria inoculation of barley against aphids is dynamic and acts through several molecular pathways to induce plant resistance (defences) and tolerance (nutrition and growth) to aphids. Future research holds promise for exploiting these interactions for sustainable crop protection and pest management in agriculture.

## Introduction

Plants encounter a broad range of insect herbivores that challenge the plant’s capability for growth, survival, and yields. In response to insect herbivory, plants activate inducible defences, including physical and chemical traits to reduce insect feeding and performance (War *et al*., 2012; Mithöfer & Boland, 2012). Plant induced resistance against insect herbivores is regulated by complex interactions among various plant secondary metabolites, phytohormone signalling, nutritional, and structural pathways (Pieterse *et al*., 2009). These pathways interact cooperatively and antagonistically, shaping the specific nature of plant defence responses to the current environment (Pieterse *et al*., 2009; Li *et al*., 2019).

Plants naturally associate with multitude of beneficial soil microbes on roots, often referred to as their second genome (Turner *et al*., 2013). These microbes, including fungi and bacteria, provide beneficial functions for the plant including nutrition provision, growth promotion, and increased resilience to biotic and abiotic stress (Pieterse *et al*., 2009; Goh *et al*., 2013; Shikano *et al*., 2017). When under attack, plants can recruit beneficial bacteria from the surrounding soil to their roots to boost their defences against herbivores (van Loon *et al*., 1998; Berendsen *et al*., 2012). The presence of beneficial microbes before herbivory can provide an added advantage, as rhizobacteria can prime plant defences that are only triggered upon herbivory (Conrath, 2009; Kim & Felton, 2013). Primed plants respond more quickly and with stronger defences responses than non-primed plants (Martínez-Medina *et al*., 2016). Microbial inoculation of plants can provide access to these beneficial microbes for plants before herbivore arrival, akin to a plant vaccination for stress-resilience (Berendsen *et al*., 2012). A recent meta-analysis highlighted the broad potential of rhizobacterial inoculation of plants roots to suppress insects on the plants, with a clear focus on crop pests (Zytynska *et al*., 2024). For piercing and sucking insects such as aphids, significant effects on reducing insect fecundity were dominated by rhizobacteria *Bacillus* strains, while the rhizobacteria *Pseudomonas* spp. and other less studied rhizobacteria showed negative effects on insect fitness, depending on the experimental system (Zytynska *et al*., 2024).

The vast majority of microbe-mediated induced resistance has primarily been studied using marker genes for plant-microbe-insect studies, focused on a central role of phytohormones. In *Arabidopsis thaliana*, inoculation with *P. fluorescens* primed the plants for enhanced expression of Lipoxygenase 2 (*LOX2*) genes involved in jasmonic acid-regulated defence pathways upon aphid *M. persicae* feeding (Pineda *et al*., 2012). However, *Bacillus velezensis* inoculation of *A. thaliana* induced higher accumulation of hydrogen peroxide, cell death, and callose deposition in leaves compared to untreated plants with no role of salicylic acid, jasmonic acid, ethylene, or abscisic acid in the defence against *Myzus persicae* aphids (Rashid *et al*., 2018). In tomato, inoculation of *B. subtilis* triggered jasmonic acid dependent as well as jasmonic acid independent responses to reduce whitefly *Bemisia tabaci* (Valenzuela-Soto *et al*., 2010). In barley, *Acidovorax radicis* was shown to induce plant flavonoids but was not tested in relation to aphid feeding (Han *et al*., 2016), while a later study again found only limited evidence for the role of salicylic acid, jasmonic acid, or ethylene with potential effects again identified for the flavonoid biosynthesis pathway (Sanchez-Mahecha *et al*., 2022).

Here, we present the first comprehensive global transcriptomic study comparing the inoculation of two distantly related rhizobacteria (*Acidovorax radicis* and *Bacillus subtilis)* individually inoculated to barley plant roots, and the effect on a piercing-sucking aphid herbivore. Given that microbe-plant-insect interactions are dynamic, and that plant responses to insect herbivory can differ depending on feeding duration and subsequent insect infestation densities, we hypothesized that the impact of the rhizobacteria inoculation and induced transcriptomic changes against the aphid is time dependent. We performed a bioassay of barley plants inoculated with rhizobacteria and uninoculated controls, and evaluated aphid fitness, by assessing aphid population growth at 7, 14 and 21 days after aphid infestation; also, with no-aphid control plants. We performed transcriptomic (RNA*seq*) analyses to elucidate plant global gene expression using a fully factorial design to disentangle effects of rhizobacteria and aphid feeding; we harvested and sequenced plants after 24 hours (initial aphid feeding), 7 days (aphid colony establishment) and 21 days (sustained aphid population growth) of aphid feeding. Our study sheds light on underlying molecular changes responsible for defence induction and priming of defences by rhizobacteria with impacts on the aboveground aphid populations.

## Material and methods

### Plants, rhizobacteria and insect-herbivores

We used barley (*Hordeum vulgare*, KWS Irina) individually inoculated with two different rhizobacteria species, *Acidovorax radicis* (N35) and *Bacillus subtilis* (B171). We used the English grain aphid *Sitobion avenae* (clonal genotype Fescue maintained as a lab population for >10 years) as the leaf feeding insect herbivore. All experiments were conducted in a plant growth chamber (Fitotron SGC120, Weiss Technik UK Ltd) set at 18 °C, 16 h:8 h light-dark photoperiod cycle, and 65 % relative humidity.

### Cultivation of rhizobacteria Acidovorax radicis and Bacillus subtilis

*Acidovorax radicis* was cultivated at 30 °C on nutrient agar plates (Difco™ General purpose nutrient broth 8 g/litre plus 15 g -agar) and grown for 96 hours. Cells were harvested and collected into 10 mM MgCl_2_ and homogenized through a 0.8 µM syringe needle. We resuspended *A. radicis* at a final optical density (OD) 600 nm of 2.0 (estimated 1.6×10^9^ CFU/ml). We grew *B. subtilis* for 24 hours at 30 °C in nutrient-broth (Difco™ General purpose Nutrient Broth 8 g/litre) while shaking at 220 rpm on Innova 2300 platform shaker (New Brunswick Scientific Europe B.V Nijmegen Netherlands). Cells were harvested by centrifuging the broth at 4 °C, and at 4000 xg for 10 minutes in Sorvall LYNX 4000 Centrifuge (Thermo Scientific, Osterode am Harz, Germany), cleaned in 0.9 % NaCl and finally suspended in 10 mM MgCl_2_, and resuspended to a final *B. subtilis* optical density (OD) 600 nm of 2.0 (estimated 1.6×10^9^ CFU/ml). The bacterial suspensions were used immediately for the bioassays.

### Plant growth condition, rhizobacteria inoculation, and aphid infestation

We used a fully factorial experimental design including three inoculation treatments (control, *A. radicis*, *B. subtilis*) and two aphid treatments (with aphids or without aphids). Before germination, barley seeds were surface-sterilised by immersion in 4 % sodium hypochlorite (bleach) solution for 1 minute, and then rinsed thoroughly under running tap water. For rhizobacteria inoculation, seeds were soaked in the suspension of *A. radicis* or *B. subtilis* for 2 hours, whereas control seeds were mock inoculated by soaking in 10 mM MgCl_2_ for 2 hours. The inoculated seeds were then germinated in seed-trays in potting substrate (Levington’s Advance F1 Compost, low nutrient). When seedlings were six days old, we measured the root and shoot lengths to select homogenous plants for the bioassays. We transplanted the seedlings into individual pots (8.5 cm height x 9 cm diameter) and immediately watered each with 60 ml of tap water, and afterwards as required (approximately every 3 days). The seedlings were grown in the plant growth chamber for 24 hours until approx. 10 cm tall before being used for the experiment. Aphid infestation was achieved by placing six *S. avenae* apterous (unwinged) 4^th^ instar aphids to the base of the plant using a fine paintbrush. Every plant pot was then covered with a fine white net, using a frame, to eliminate aphid movement across plants. Further we placed the plant pots onto individual saucers to avoid soil or rhizobacteria cross-contamination. Each treatment was replicated six times, for each harvesting timepoint of 24 hours, 7 and 21days after aphids feeding. We recorded plant shoot lengths and counted the number of aphid adults and offspring to estimate the level of herbivory. For 21 days experiment plants, we counted the number of aphids at day 7, 14 and 21 after aphid infestation to evaluate aphid population growth over time. Shoot tissue was collected and flash-frozen in liquid nitrogen and stored at −80 °C freezer.

### RNA extraction and sequencing

We extracted total RNA from leaves samples collected at the three study timepoints (24 hours, 7 days, 21 days). We used 50 ± 5 mg (fresh weight) of ground leaf material and extracted the RNA using TRizol reagent (Thermo Fisher Scientific, United Kingdom), according to the manufacturer’s instructions. All samples were treated with 2U/ µl of RNAse-free DNase I (Thermo Scientific Fisher, United Kingdom) to remove any traces of DNA. The RNA quantity and quality were confirmed using ND-1000 UV/vis spectrophotometer (NanoDrop Technologies, Wilmington, Delaware, USA), and by gel electrophoresis (1 % molecular grade agarose). A Qubit 4 Fluorometer (Life Technologies Holdings Pte Ltd, Woodland HDB Town, Singapore) was used to assess the RNA integrity. Samples whose RNA integrity quality score was ≥ 7 were selected for library preparation and sequencing at the Centre for Genomic Research (CGR), University of Liverpool in the UK. We sequenced four replicates from every treatment combination (72 samples). The preparation of RNA libraries involved depletion of Ribosomal-RNA via poly-A selection using NEBnext Poly(A)mRNA magnetic isolation module. The poly A enriched samples were then used as input in the NEBnext Ultra II RNA kit (NEB, USA), and processed following the manufacturer’s instructions. Libraries were validated and quantified using the Illumina library quantification kit (KAPA, Library Quant Kit (Illumina) *q*PCR master mix 2x) on Roche Light Cycler LC480II (Roche Diagnostic Ltd, Switzerland) following manufacturer’s recommendations. Libraries were sequenced on an Illumina NovaSeq 6000 platform (Illumina, San Diego, CA, USA) following standard workflow over 1 lane of an S4 flow cell. The sequencing produced 2 x 150 bp paired-end raw reads with a mean sequencing depth of ∼40 million reads per sample. Raw read sequence files will be available on the Gene Expression Omnibus (GEO; accession number [upon publication]).

### Transcriptomic analysis

The raw Illumina reads were first pre-processed for initial quality assessment using the in-house pipeline at the CGR-genomics at the University of Liverpool. Base calling and de-multiplexing of indexed reads were carried out using CASAVA v.1.8.2 conversion software (Illumina) to produce read sequences in fastq format. Clean reads were obtained by using Cutadapt v.1.2.1 software by trimming the Illumina adapter sequences on the raw fastq files (Martin, 2011). We removed low-quality bases using Sickle v.1.200 software set at minimum window quality score 20. Raw reads shorter than 20 bp were removed from the dataset. The clean high-quality paired-end raw reads were pseudo-aligned to the barley reference transcriptome sequence (Ensemblplants; Hordeum vulgare.MorexV3_pseudomolecules_assembly.cdna.all.fa) using Kallisto v. 0.46.2 (Bray *et al*., 2016). The number of raw reads mapped to each gene was counted using the kallisto quant function.

### Differential gene expression and functional enrichment analyses

We used statistical computing environment R version 4.3.2 in RStudio version 2022.12.0+353 and Bioconductor version 3.16 for differential gene expression and functional enrichment (Gentleman *et al*., 2004; Huber *et al*., 2015; R Core Team, 2022; RStudio-Team, 2022). First, we summarized the reads (transcripts) count data at the gene level using the tximport R package (Soneson *et al*., 2016). Then normalized it using the trimmed mean of M values (TMM) method in edgeR (Robinson *et al*., 2010). We filtered genes with less than 1 count per million (CPM) in n+1 (where n is the number of biological replicates in treatment group). The single and interactive effects of rhizobacteria and aphid on gene expression were determined using PERMANOVA models in R including rhizobacteria treatment (control, *A. radicis* or *B. subtilis*), aphids (*S. avenae*) and their interaction as explanatory variables. Then we used the voom function in limma R package for variance stabilization (Ritchie *et al*., 2015). The limma R package and decideTests function was used to select differentially expressed genes at FDR <0.05, absolute logFC ≥ +2 or ≤ −2 for each study timepoint (Benjamini & Hochberg, 1995). Gene ontology (GO) enrichment analyses were performed using a self-curated annotation file based on the *Hordeum vulgare* gene (MorexV3_pseudomolecules_assembly) dataset in the Ensemblplants public database. We extracted the TERM2NAME and TERM2GENE mapping from the GO.db R package (Carlson, 2022) and used it for GO enrichment over-representation analysis (ORA). Significantly enriched GO categories, including biological process (BP), molecular functions (MF) or cellular components (CC) terms, were determined at p-value Cutoff ≤ 0.05, P-value adjusted by Benjamin Hochberg method, and q-value Cutoff ≤ 0.2. KEGG enrichment analyses was achieved by converting the differentially expressed genes into rice orthologs (*Oryza sativa* Japonica group) using g:Profiler2 the web application (Kolberg *et al*., 2020). This approach was ideal because barley is not yet fully annotated and lacks enrichment KEGG datasets. The rice Orthologs were enriched in the rice KEGG database RAPDB. Significantly enriched KEGG pathways were determined at p-value Cutoff ≤ 0.2, P-values were adjusted by the Benjamin Hochberg method, and q-value Cutoff ≤ 0.2 (Li, 2022).

### Gene co-expression analyses

We performed weighted gene co-expression network analyses (WGCNA) using the WGCNA R package to detect co-expressed gene modules related to the impact of rhizobacteria against aphids (Langfelder & Horvath, 2008). We used raw read counts data and filtered out genes with less than 50 counts across all treatment per study timepoint. Then we normalised the data and calculated the power value for gene correlation using the pickSoftThreshold function (Fig. S1a,c,e). We used the block-wise modules function to construct block-wise networks with the following parameters: power = selected power from fitIndices, networkType = signed, maxBlockSize = 5000, TOMType = signed, minModuleSize = 30, mergeCutHeight = 0.25, and plotted dendrograms of the identified gene modules (Fig. S1b,d,f). Next, we assessed the module-trait relationship by performing linear models in R (limma) to calculate eigengene (hypothetical central genes) for each module, and extracted the gene module exhibiting largest significant difference across treatment groups. Further, we extracted genes in the selected modules of interest and performed functional enrichment in KEGG database as described above for differentially expressed genes. Gene connectivity based on edge weights (ranging 0 to 1) was determined by topological overlap matrix (TOM). The weight across all edges of a node were summed and used to define the level of connectivity. Nodes with high connectivity (≥100) were presented in the network. Module-membership or intramodular connectivity measure (connectivity of a gene to all other genes) within same module was used to define hub genes (driver genes) in each module per timepoint; module eigengene-based connectivity (kME) values ≥ 0.8 were considered as the hub genes for the particular module. Similarly, we performed functional enrichment of the hub genes in the KEGG database as described above.

## Results

### Rhizobacteria suppress aphid populations on barley

We first examined the effect of rhizobacteria inoculation on performance of the aphid. Overall, our results demonstrated significant suppression of the aphid *S. avenae* population on plants inoculated with either rhizobacteria (*A. radicis* or *B. subtilis)* compared to control plants, and across the three timepoints (7 days: F_2,11_=8.47, P=0.018; 14 days: F_2,11_=12.98, P=0.007; and 21 days: F_2,11_=6.93, P=0.028, Fig. **1**).

**Fig. 1.**
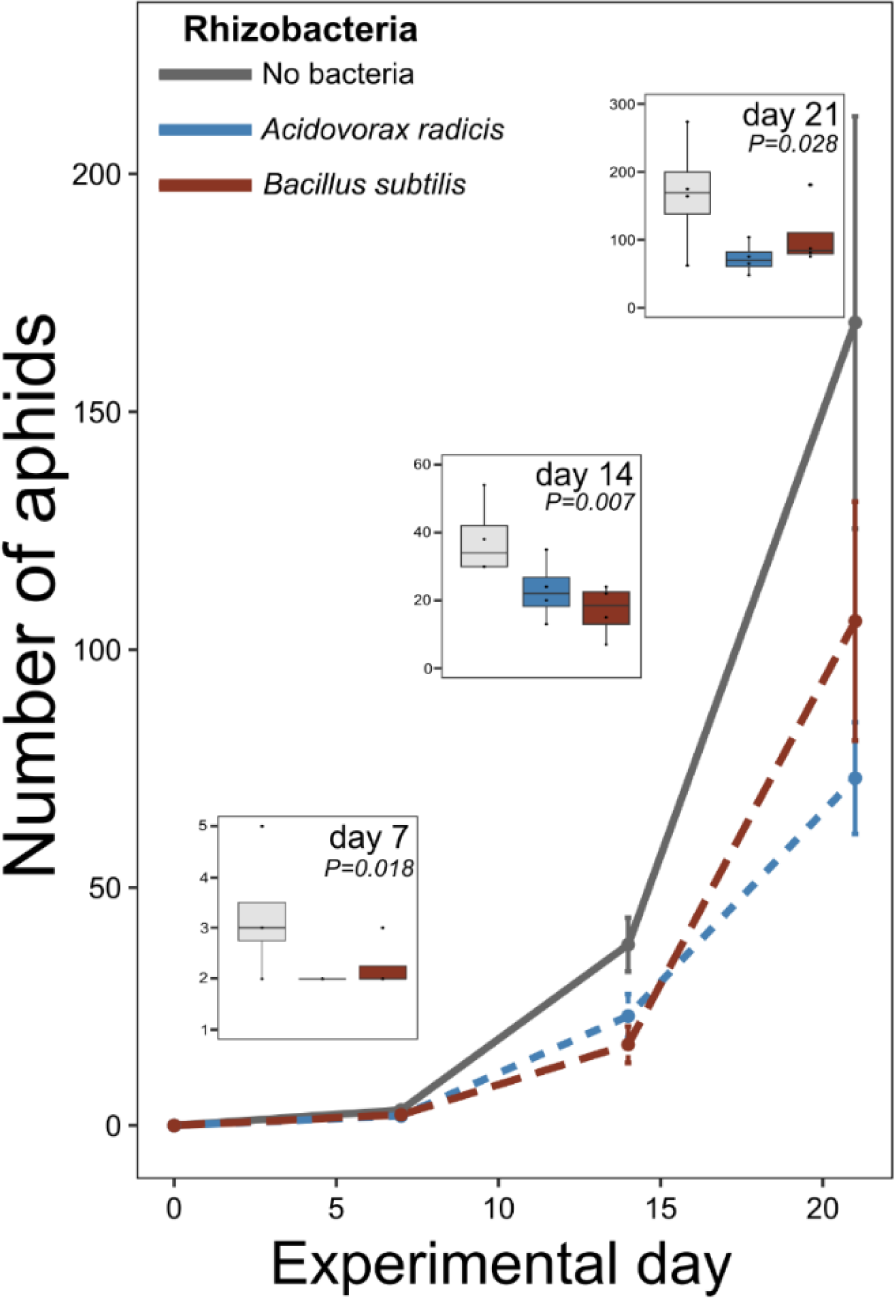
Impact of plant inoculation with rhizobacteria *Acidovorax radicis* or *Bacillus subtilis* on the population growth of the aphid *Sitobion avenae*. Insert plots show data within time point with associated statistical significance, line plot shows this across the experimental time. Error bars are mean ±standard error (*n* = 4).

### Rhizobacteria and aphids trigger distinct transcriptome profiles in barley

To unravel the plant responses altered by rhizobacteria resulting in suppression of the aphid populations, we performed RNA*seq* analyses at three timepoints, including 24 hours (initial aphid feeding), 7 days (aphid colony establishment) and 21 days (sustained aphid population growth) after aphid infestation. We constructed 72 RNA libraries, and obtained an average of 38M reads per library. We mapped 74 % of the clean raw reads to the barley reference transcriptome (Table **S1**), and identified about 24,000 annotated genes. We found that the modulation of the barley transcriptome by rhizobacteria or aphids differed significantly across the timepoints (PERMANOVA: F_2,63_=31.15, R^2^=0.46, P<0.001). We detected significant main effects of aphid feeding (averaged across timepoints: F_1,63_=4.35, R^2^=0.03, P=0.002) and rhizobacteria root inoculation dependent on time (interaction between bacteria and timepoint: F_2,63_=2.25, R^2^=0.03, P=0.012). Because our experiment involved two different bacterial species, we checked for species specific patterns. However, we found that rhizobacteria-induced changes in barley global transcriptomic profile were not specific to the bacteria strain (main effect of strain identity: F_1,40_=0.63, R^2^=0.01, P=0.711). Our results suggest that the observed changes in barley transcriptome are explained by interactive effects across our treatments. At 24 hours after aphid feeding, we observed strong significant interaction effects (aphid-by-rhizobacteria presence) on transcriptomic changes (Fig. **2a**; Table **1**). After 7 and 21 days of aphid feeding, the most significant factor modulating plant transcriptomic changes were the aphids, with decreasing strength of effects of rhizobacteria inoculation (Fig. **2d,g**; Table **1**).

**Fig. 2.**
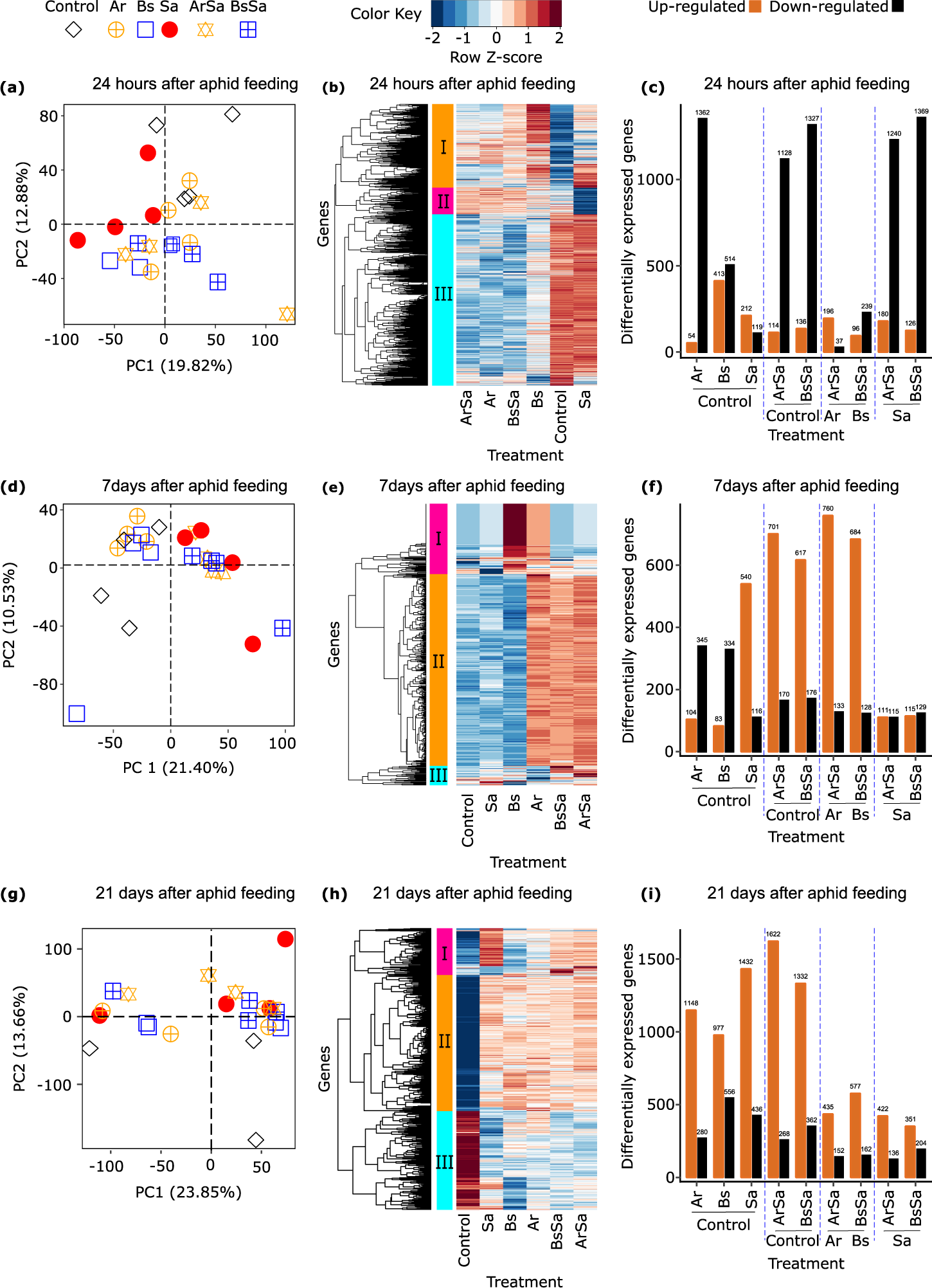
Induced transcriptome profiles of barley leaves upon rhizobacteria root colonization and shoot herbivory. Overview of the global transcriptome profile and differentially expressed (DE) genes in leaves of barley plants without treatment *(Control),* inoculated with rhizobacteria *Acidovorax radicis* (Ar) or *Bacillus subtilis* (Bs), infested with aphid *Sitobion avenae* (Sa), or inoculated with rhizobacteria and infested with aphids (ArSa or BsSa). The panels are arranged per time point based on aphid feeding duration, including 24 hours in panels **a-c**, 7 days in panels **d-f**, and 21 days in panels **g-i**. Panels **a,d,g** in the first column are PCAs showing scores, panels **b,e,h** in the middle column are heatmaps of DE genes across treatment (each line or row represents a unique gene, and each column represents a unique treatment group). Panels **c,f,I** in the third column are bar plots showing the number of DE genes per treatment relative to the respective control signified by different colours (blue indicates downregulation, white indicates no change, while red indicates upregulation).

**Table 1.** Permutation Multivariate Analyses of Variance (PERMANOVA) results of transcriptome expression upon rhizobacteria root colonization and shoot herbivory.

| Source of variation | Time points |  |  |  |  |  |  |  |  |  |  |  |
| --- | --- | --- | --- | --- | --- | --- | --- | --- | --- | --- | --- | --- |
|  | 24 hours |  |  |  | 7 days |  |  |  | 21 days |  |  |  |
|  | df | F | R2 | P | df | F | R2 | P | df | F | R2 | P |
| Rhizobact.P/A | 1,19 | **4.17** | **0.16** | **<0.001** | 1,20 | 1.47 | 0.06 | 0.074 | 1,19 | 1.42 | 0.06 | 0.104 |
| Aphid | 1,19 | 1.16 | 0.04 | 0.245 | 1,20 | **4.48** | **0.17** | **<0.001** | 1,19 | **2.03** | **0.09** | **0.014** |
| Rhizobact.P/A*Aph | 1,19 | **2.19** | **0.08** | **0.007** | 1,20 | 0.83 | 0.03 | 0.693 | 1,19 | 1.23 | 0.05 | 0.187 |

§ Rhizobact.P/A, rhizobacteria presence/ absence; Aph, aphid; df, degrees of freedom P-values in bold indicates statistical significance with $P < 0.05$

Next, we obtained differentially expressed genes and performed hierarchical clustering, revealing three main gene clusters varying across the treatments and timepoints (Fig. **2b,e,h**). After 24 hours of aphid feeding, most up-regulated genes were contained in Cluster III from control and aphid-infested plants, while this cluster was predominantly down-regulated in plants inoculated with rhizobacteria regardless of aphid presence (Fig. **2b**). After 7 days of sustained aphid feeding, Cluster II contained majority of the differentially-expressed genes. These genes were highly expressed in *A. radicis* inoculated plants with and without aphid infestation, and in *B. subtilis* inoculated plants only when aphids were present (Fig. **2e**). After 21 days of aphid feeding and population growth, there was a notable separation of control plants to all other treatments, with less variation in gene expression among all inoculated plants than in the other timepoints (Fig. **2h**). To further examine the interactive effects, we compared the different treatments relative to their respective controls. Gene expression increased over time, with the combined (interactive) effect of rhizobacteria inoculation and aphid infestation driving the majority of gene expression differences (Fig. **2c,f,i**; Table **S2**). At day 7 when aphid colonies were establishing, we observed a strong induction of aphid-related genes (Fig. **2f** *Sa vs control* and *ArSa vs Ar*, *BsSa vs Bs;* Table **S2**).

Using KEGG pathway enrichment analyses we showed that the differentially expressed genes were enriched in 16 KEGG subcategories belonging to five main categories namely, metabolism, genetic information processing, environmental information processing, organismal systems and cellular process (Fig. **3**). Notably, our transcriptome data showed that metabolism was most affected especially the glutathione (amino acid) and phenylpropanoid (secondary metabolite) pathway. The main modulated glutathione gene was the *glutathione-S-transferase* (GST). While several phenylpropanoid genes were altered, including *phenylalanine-ammonia lyase* (PAL), *Caffeic acid -O-methyltransferase* (CaOMT), *Cinnamoyl-CoA reductase* (CCR), *Flavonoid 3’-O-methyltransferase, Sinaply-alcohol dehydrogenase* (SAD). In addition to these phenylpropanoid genes, we found modulation of *peroxidases* highlighting their involvement in regulating the phenylpropanoid pathway (Table **S3-S6**).

**Fig. 3.**
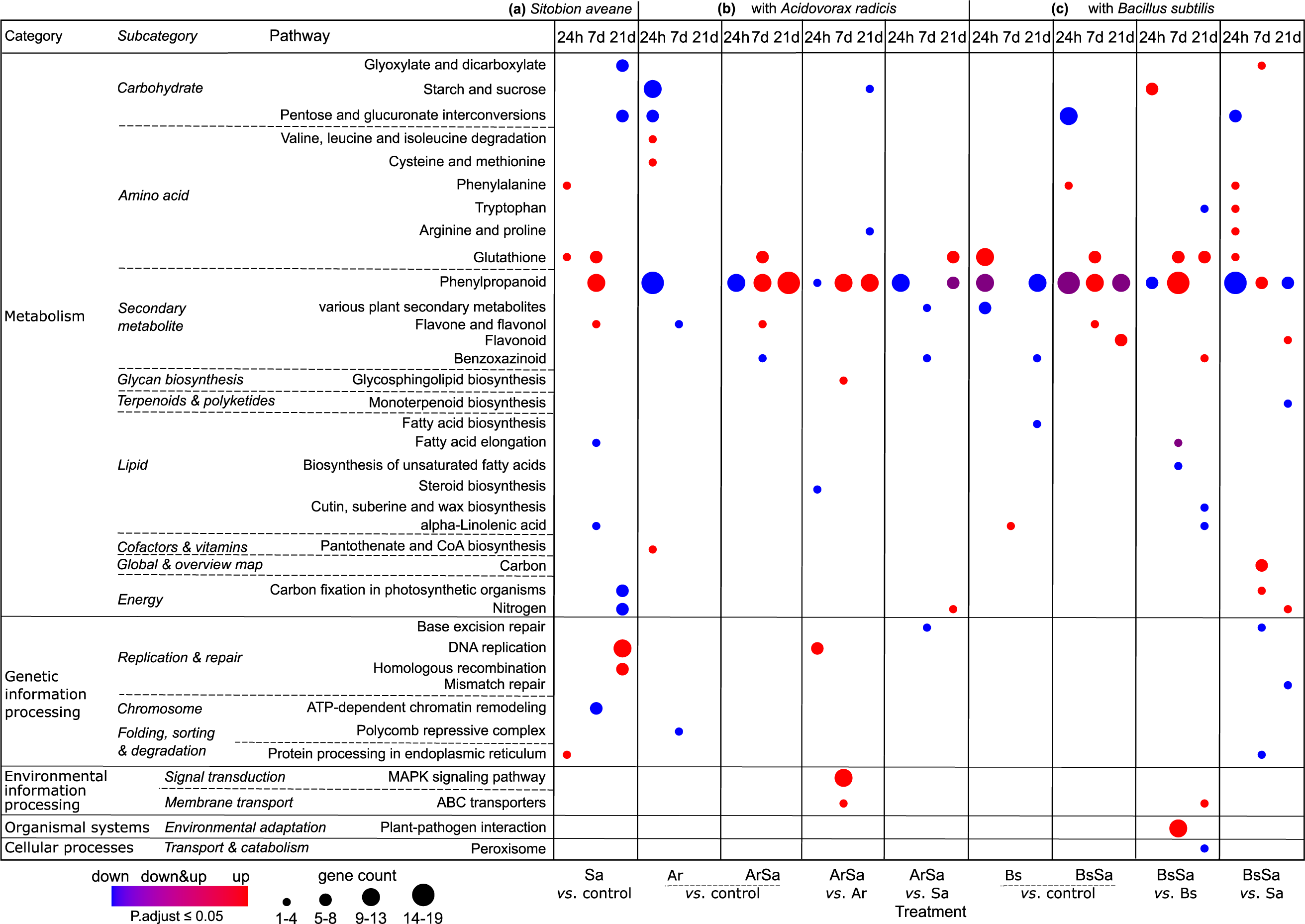
KEGG pathway enrichment analysis of differentially expressed genes in barley leaves upon rhizobacteria root colonization and aphid herbivory. This bubble plot showed the enriched KEGG pathways of upregulated and downregulated differentially expressed genes in leaves of barley without treatment *(Control),* infested with the aphid *Sitobion avenae (Sa),* inoculated with the rhizobacteria *Acidovorax radicis (Ar)* or *Bacillus subtilis (Bs)*, or inoculated with rhizobacteria and infested with aphid *(ArSa* or *BsSa)*. In *Sa*, *ArSa* and *BsSa* plants, the aphid *S. avenae* was allowed to feed on plants for 24 hours, 7 and 21 days. **(a)** shows genes induced by *Sa* relative to *Control*. **(b)** shows treatment groups inoculated with *Ar*: *Ar vs. Control* are genes altered by *Ar* in absence of aphids. *ArSa vs. Control* are genes induced by both rhizobacteria and aphids relative to *Control*. *ArSa vs. Ar* shows the aphid *(Sa)* induced genes in rhizobacteria *(Ar)* inoculated plants. *ArSa vs. Sa* shows the rhizobacteria *(Ar)* induced genes in aphid *(Sa)* infested plants. **(c)** shows treatment groups inoculated with *Bs*: *Bs vs. Control* are genes altered by *Bs* in absence of aphid. *BsSa vs. Control* are genes induced by both rhizobacteria *(Bs)* and aphid *(Sa)* relative to *Control*. *BsSa vs. Bs* shows the aphid *(Sa)* induced genes in rhizobacteria *(Bs)* inoculated plants. *BsSa vs. Sa* shows the rhizobacteria *(Bs)* induced genes in aphid *(Sa)* infested plants.

Overall, aphid infestation of plants without rhizobacteria inoculation up-regulated genes in the glutathione pathway at 24 hours and 7 days, and phenylpropanoid pathway after 7 days of feeding (Fig. **3**, *Sa vs. Control*). In rhizobacteria-inoculated plants without aphid infestation, *B. subtilis* induced glutathione at the 24 hr timepoint (Fig. **3**, *Bs vs. Control*), while there were genes both up- and down-regulated in the phenylpropanoid pathway indicating dynamic expression in this pathway (Fig. **3**, *Bs vs. Control*). For *A. radicis* inoculated plants (no aphids), there was strong suppression of the phenylpropanoid pathway at the 24 hr timepoint, with no induction of the glutathione pathway (Fig. **3**, *Ar vs. Control*). With aphid feeding, we still observed *A. radicis* suppression of the phenylpropanoid pathway at the 24 hr timepoint but also further suppression effects at 21 days (Fig. **3**, *ArSa vs. Sa*). Aphid induction of phenylpropanoid defences continued through to 21 days on inoculated plants (Fig. **3**, *ArSa vs. Ar*), which was not observed on uninoculated plants (Fig. **3**, *Sa vs. Control*). A similar pattern was observed for *B. subtilis* plants, where the rhizobacteria suppressed the phenylpropanoid pathway at the 24 hr and 21 day timepoints (Fig. **3**, *BsSa vs. Sa)*, potentially mitigating some of the aphid-induction of these defences (Fig. **3**, *BsSa vs. Bs)*.

A number of other pathways across the main categories showed variable regulation across our treatments. After 21 days of aphid feeding in absence of rhizobacteria, aphids down-regulated carbohydrates, including *glyoxylate and dicarboxylate* and *pentose and glucoronate interconversion*. Starch and sucrose carbohydrate pathways, such as *beta 1-4 glucanase* and *α-glucoside,* were down-regulated by *A. radicis* inoculation at the early timepoint (Fig. **3**, *Ar vs. Control)*, while *B. subtilis* inoculation only suppressed these upon aphid feeding (Fig. **3**, *BsSa vs Sa)*. *Bacillus subtilis* further induced expression of genes for carbohydrate-related *Glyoxylate* and *dicarboxylate*, including *ribulose 1, 5-bisphosphate carboxylase* at day 7 (Fig. **3**, *BsSa vs Sa*), which were suppressed by aphids on control plants in the later time point (Fig. 3 *Sa* vs *Control*). We also found up-regulation of *replication* and *repair genes* at day 21 that may reflect plant responses to increased aphid population sizes and subsequent damage (Fig. **3**, *Sa vs. Control*).

### Gene co-expression reveal rhizobacteria prime defence and nutritional responses in barley to suppress aphids

To identify groups of genes with similar functions or involved in same pathway that respond to rhizobacteria inoculation and/or aphid feeding, we performed weighted gene co-expression network analyses. We identified 26, 38, and 23 gene modules across the three timepoints, containing between 31-9750 genes (Table **S7**). The co-expressed modules were randomly colour-coded, and we present four that showed large differences across the treatments and timepoints (Fig. **4**, Table **S8**). After 24 hours of aphid feeding, genes in the module brown (Fig. **4a**) were upregulated in plants inoculated with rhizobacteria with and without aphids feeding, and down-regulated in uninoculated plants with and without aphids (MEbrown *n* = 2555, F = 4.61, P = 0.018; Fig. **5a**, Table **S8a**). After 7 days of aphid feeding, genes in the modules turquoise and violet (Fig. **4b**) were up-regulated in all plants infested with aphids regardless of rhizobacteria inoculation (MEturquoise *n* = 3353, F = 4.03, P = 0.022, and MEviolet *n* = 90, F = 4.12, P = 0.022; Fig. **5b**, Table **S8b**). After 21 days of aphid feeding, genes in the module grey60 (Fig. **4c**) were upregulated across the aphid infested plants, while *A. radicis* inoculated plants without aphids showed intermediate expression (MEgrey60 *n* = 85, F = 3.40, P = 0.059; Fig. **5c**, Table **S8c**).

**Fig. 4.**
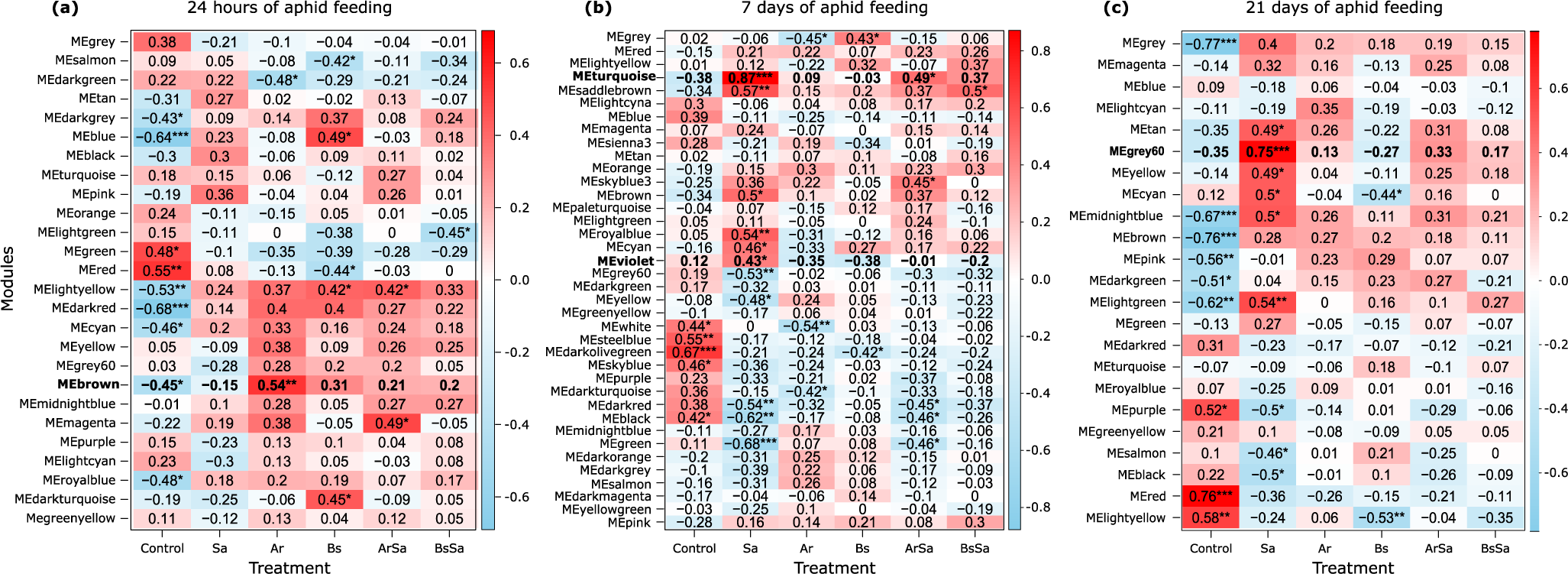
Overview of gene co-expression modules in barley leaves upon rhizobacteria root colonization and shoot herbivory. Panels are heatmaps showing the relationship between each gene co-expression module and the treatment groups. The treatment groups include barley plants without treatment *(Control),* infested with the aphid *Sitobion avenae (Sa),* inoculated with the rhizobacteria *Acidovorax radicis (Ar)* or *Bacillus subtilis (Bs)*, or inoculated with rhizobacteria and infested with aphid *(ArSa* or *BsSa)*. In *Sa*, *ArSa* and *BsSa* plants, the aphid *S. avenae* was allowed to feed on plants for 24 hours **(a)**, 7 days **(b)** and 21 days **(c)**. Each row represents a module named randomly by colour. Each column represents a treatment group. The colour of each cell represents the association coefficient between module and the treatment group, red colour denotes positive while blue indicates negative relationship between the module and treatment. Asterisks indicate a significant correlation.

**Fig. 5.**
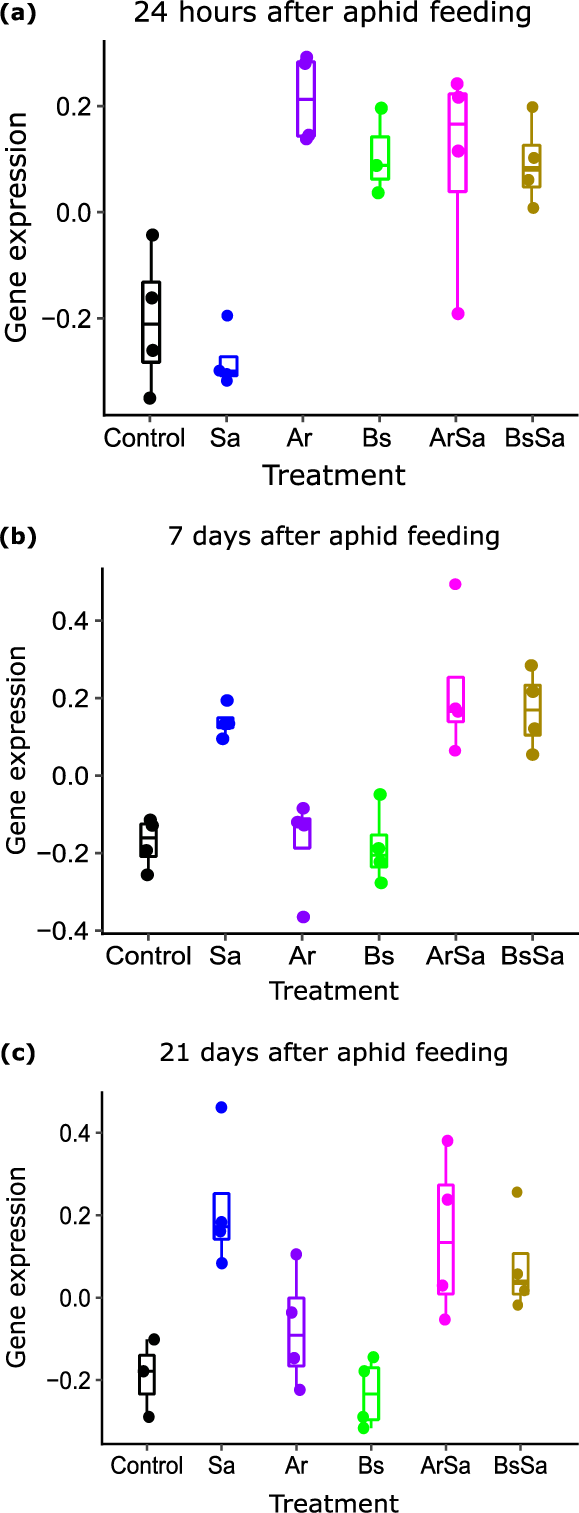
Gene co-expression pattern of selected modules of interest over time points in barley leaves upon rhizobacteria root colonization and shoot herbivory. **(a)** brown module at 24 hours, **(b)** combined turquoise and violet modules at 7 days, and **(c)** grey60 module at 21 days post aphid infestation. The treatment groups include barley plants without treatment *(Control),* infested with the aphid *Sitobion avenae (Sa),* inoculated with the rhizobacteria *Acidovorax radicis (Ar)* or *Bacillus subtilis (Bs)*, or inoculated with the rhizobacteria and infested with the aphid *(ArSa* or *BsSa)*.

Next, we performed KEGG enrichment analyses of the genes in the selected modules of interest at each timepoint. After 24 hours of aphid feeding, we found co-expressed genes (‘brown’ module) enriched for plant hormone signal transduction, phenylpropanoid biosynthesis, and arachidonic acid metabolism (Fig. **6a**). In the plant-hormone signal transduction, we found the co-expressed genes included, *ubiquitin ligase complex*, *jasmonyl-L-isoleucine synthase* for JA biosynthesis, *ethylene receptor and response* for ET signaling, *auxin responsive protein, auxin transporter, indole-3-acetic acid protein* for AUX responses, *histidine phosphotransfer protein* for CK signaling, and *mitogen-activated protein kinase kinase 4* for activation of the MAPK (Table **2**, Table **S9a**). For the phenylpropanoid pathway, we found seven main genes, including *phenylalanine-ammonia lyase* (PAL), *4-coumarate-CoA*, *cinnamyl alcohol dehydrogenase* (CAD), *sinapyl alcohol dehydrogenase* (SAD), *cinnamoyl-CoA reductase* (CCR), *caffeic acid O-methyltransferase*, *flavonoid 3’-O-methyltransferase* that lead to lignin and flavonoid biosynthesis. In addition, the phenylpropanoid genes were co-expressed with several *peroxidases* suggesting they play active role in regulating the phenylpropanoid pathway (Table **2**, Table **S9a**). In the arachidonic acid metabolism pathway, we found five genes, among them a gene similar to *gamma-glutamyl transpeptidase 4* which suggest regulatory link between arachidonic acid metabolism and glutathione pathway (Table **2**, Table **S9a**). After 7 days of aphid feeding the co-expressed genes (modules ‘turquoise’ and ‘violet’) were enriched for amino sugar and nucleotide sugar metabolism and biosynthesis of nucleotide sugar (Fig. **6b**). Among some of the main sugar genes includes *hexokinase*, *UDP-glucose dehydrogenase*, *mannose-1-phosphate guanyl transferase*, and *fructokinase*. These gene were co-expressed with *chitinases*, including *chitinase 4, 5* and *8* (*pathogenesis-related (PR)-3*) suggesting activation of constitutive mechanisms in addition to the nutritional responses (Table **2**, Table **S9b**). While 21 days after aphid feeding (‘grey60’ module), we observed co-expression among genes encoding for flavone and flavonol biosynthesis pathways which is a branch of phenylpropanoid biosynthesis. These *UDP-glucuronosyl/UDP-glucuronosyltransferase* genes were co-expressed with genes regulating biosynthesis of amino acid metabolic pathways such as *phenylalanine* and *tyrosine* that are substrates in the phenylpropanoid pathway (Fig. **6c**; Table **2**, Table **S9cd**).

**Fig. 6.**
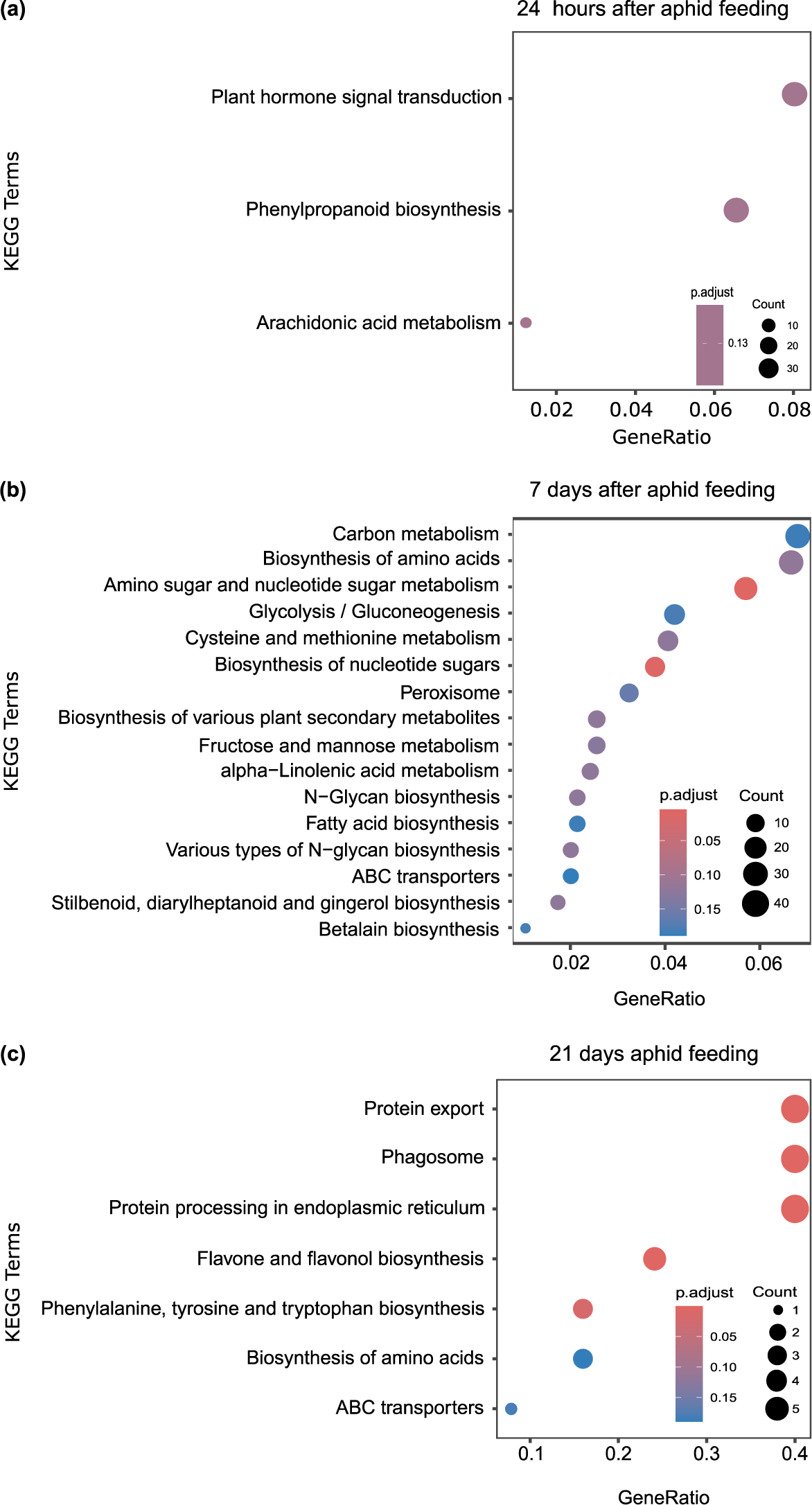
KEGG pathway enrichment analysis of co-expressed gene module in barley leaves upon rhizobacteria root colonization and shoot herbivory. Panels represent the enriched KEGG pathways of the genes in each selected co-expressed modules across all treatment groups, presented as bubble plots. The treatment groups were barley plants without treatment *(Control),* infested with the aphid *Sitobion avenae (Sa),* inoculated with the rhizobacteria *Acidovorax radicis (Ar)* or *Bacillus subtilis (Bs)*, or inoculated with rhizobacteria and infested with the aphid *(ArSa* or *BsSa)*. In *Sa*, *ArSa* and *BsSa* plants, the aphid *S. avenae* was allowed to feed on plants for (**a**) 24 hours, (**b**) 7 days and (**c**) 21 days.

**Table 2.** Enriched KEGG pathways of the co-expressed gene from selected modules in barley leaves upon rhizobacteria root colonization and shoot herbivory.

| Time points | Module | KEGG pathway | Gene count | P-value |
| --- | --- | --- | --- | --- |
| 24 hours | brown | Arachidonic acid metabolism | 5 | 0.124 |
|  |  | Phenylpropanoid biosynthesis (Lignin and Flavonoids) | 27 |  |
|  |  | Plant-hormone signal transduction (Jasmonic acid, Ethylene, Auxin, Cytokinin, MAPK) | 33 |  |
| 7 days | turquoise and violet | Amino sugar and nucleotide sugar metabolism | 37 | 0.004 |
|  |  | Biosynthesis of nucleotide sugars | 23 | 0.011 |
| 21 days | grey60 | Flavone and flavonol biosynthesis | 3 | <0.001 |
|  |  | Biosynthesis of amino acids (Phenylalanine, tyrosine and tryptophan biosynthesis) | 2 | 0.016 |

Gene co-expression analyses was performed on RNAseq data from barley plants root colonized by rhizobacteria *Acidovorax radialis* or *Bacillus subtilis* and followed by infestation with the aphid *Sitobion avenae* at three different time point, including 24 hours, 7 and 21 days after the aphid feeding. The table shows the selected modules and the KEGG pathways enriched in the particular module, number of genes involved and the Pvalues.

### Hub genes analysis revealed that rhizobacteria induce defense and nutritional response to suppress aphids on barley

We constructed gene co-expression networks based on weighting values and degree of genes in the selected modules of interest per timepoints (Fig. **7**). In each selected module of interest per timepoint, we identified the hub genes, including 339 in module brown at 24 hours, 906 in module turquoise and violet at 7 days and 121 in module grey60 at 21 days (Table **S10**). KEGG enrichment analyses for each set of hub genes revealed consistent results to our previous enrichment analyses on differential expressed genes and co-expressed gene modules. After 24 hours of aphid feeding, the hub genes in the module of interest (module brown, correlated with rhizobacteria inoculation effects) were enriched in the pathways; plant hormone signal transduction, fatty acid elongation and MAPK signaling pathway. Interestingly, we found hub gene associated with photosynthesis denoting plant growth (Fig. **7a**, Table **3**, Table **S11a**). After 7 days of aphid feeding, only hub genes from the module turquoise (correlated with aphid feeding) were enriched, and this was in amino sugar and nucleotide sugar metabolism (Fig. **7b**, Table **3**, Table **S11b**). After 21 days of aphid feeding, the hub genes in module grey60 (correlated with aphid feeding and *A. radicis* inoculation) were enriched in the pathway flavone and flavonol biosynthesis, and phenylalanine, tyrosine and tryptophan biosynthesis (Fig. **7c**, Table **3**, Table **S11c**).

**Fig 7.**
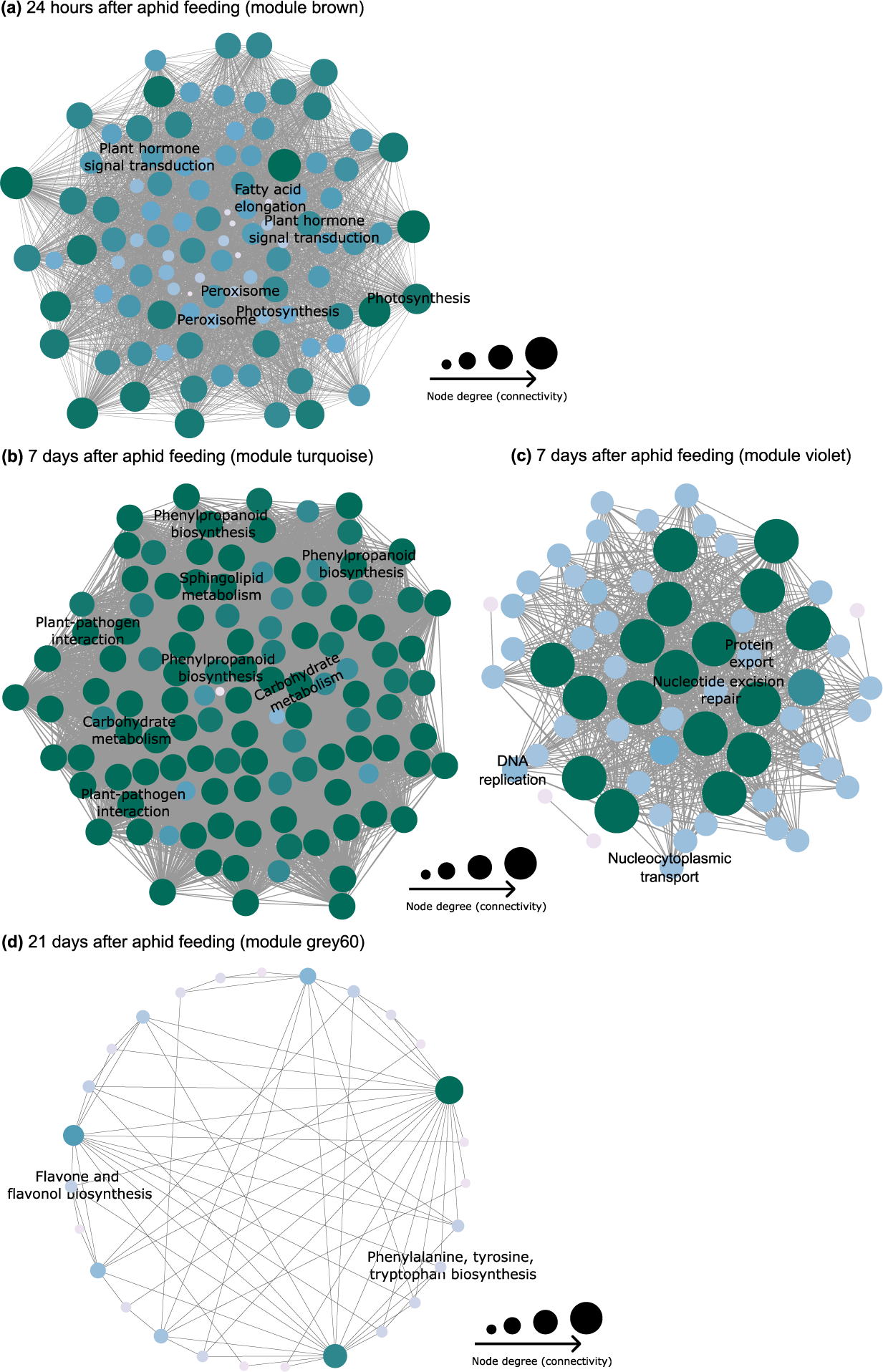
Network highlighting enriched KEGG pathways from co-expressed genes in barley leaves upon rhizobacteria root colonization and shoot herbivory. Weighted gene co-expression network analysis (WGCNA) was used to identify key modules and hub genes associated with the treatment groups. The treatment groups were barley plants without treatment *(Control),* infested with the aphid *Sitobion avenae (Sa),* inoculated with the rhizobacteria *Acidovorax radicis (Ar)* or *Bacillus subtilis (Bs)*, or inoculated with rhizobacteria and infested with aphid *(ArSa* or *BsSa)*. In *Sa*, *ArSa* and *BsSa* plants, the aphid *S. avenae* was allowed to feed on plants for (a) 24 hours (brown module), (b) 7 days (turquoise module), (c) 7 days (violet module) and (d) 21 days (grey60 module). Each node represents a gene and the connecting lines (edges) between genes represents the co-expression correlation. The node size and color are proportional to the degree of the input genes. The nodes in dark green suggest highly connected genes while the lightly coloured nodes represent less connected genes. The hub genes are labelled with the corresponding KEGG pathway term.

**Table 3.**
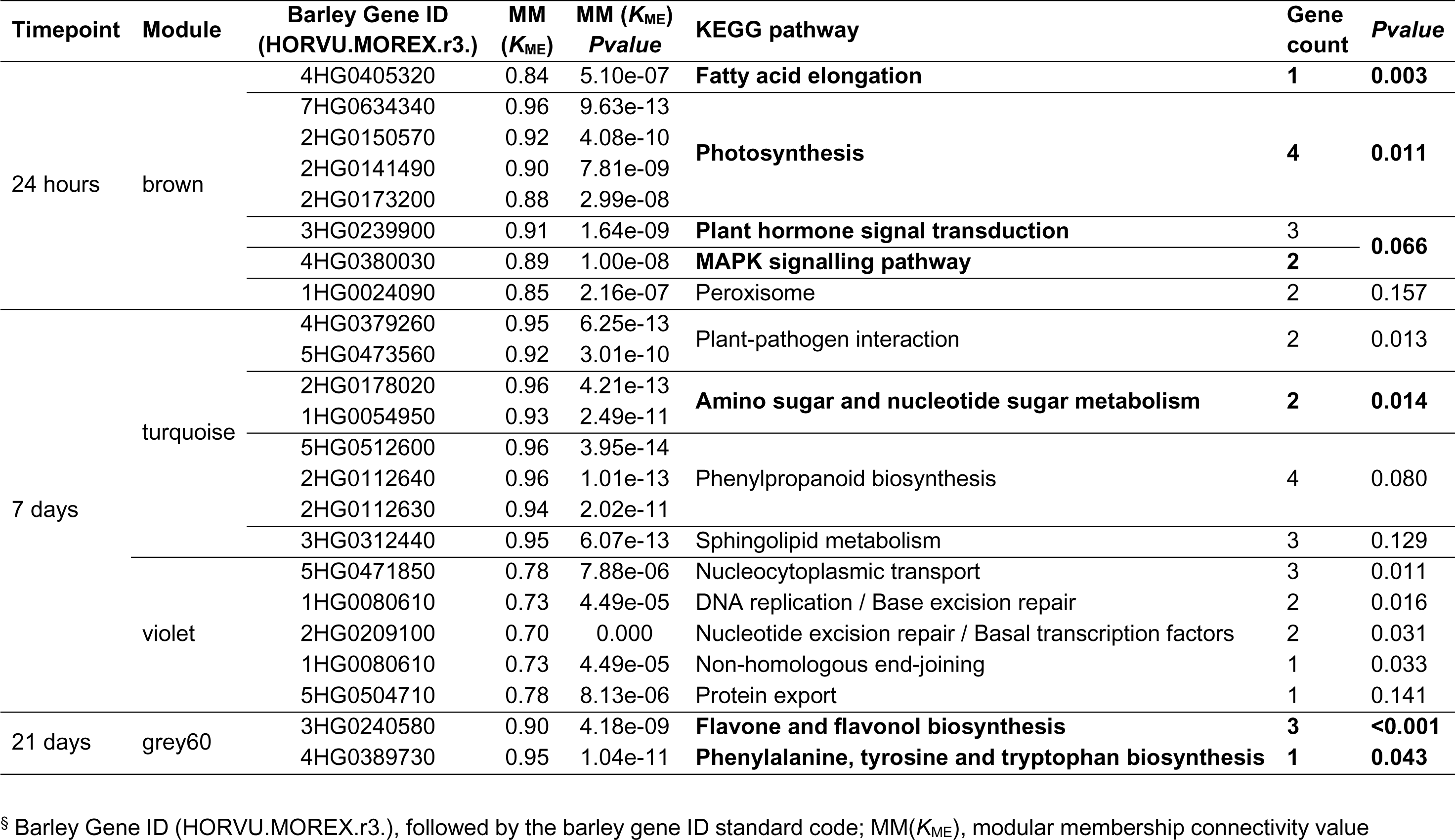
Enriched KEGG pathways of the hub genes from selected modules of co-expressed genes in barley leaves upon rhizobacteria root colonization and shoot herbivory.

## Discussion

In this study, we explored transcriptomic changes in barley inoculated with the rhizobacteria *A. radicis* or *B. subtilis*, correlated with suppression of aphid population growth on the plants. Rhizobacteria inoculation and aphids individually induced different plant responses, however it was the combined and interactive effects that drove global gene expression. After only 24 hours of aphids feeding, inoculated plants responded differently to aphids than uninoculated plants, particularly related to responses in the phytohormone, glutathione, and phenylpropanoid pathways. As aphids began to reproduce, by day 7 we observed stronger aphid-induced plant responses related to induction of the phenylpropanoid pathway but also nutrient pathways for carbohydrates, amino-acids and sugars. By day 21, aphid colonies were well established, and we observed further changes in plant defence pathways, including flavone and flavonoids, as well as genes related to plant responses to tissue damage and repair.

During early stages of aphid feeding, many of the defensive genes were suppressed on inoculated plants compared to uninoculated plants. It is possible that that rhizobacteria modulate plant defence responses to early aphid feeding when only a few aphids are present, thus conserving or reallocating energy for plant growth. This may explain our previous work on this system showing that plant growth promotion was strongest in the early growth stages of the plant (Zytynska *et al*., 2020; Xi *et al*., 2024), and also how diverse strains of rhizobacteria can simultaneously promote growth and induce resistance to herbivores (Rashid *et al*., 2018). We further identified hub genes encoding for photosynthesis during early aphid feeding supporting that rhizobacteria inoculation led to improved plant growth. Aphid-induction of defence and nutritional pathways at the 7-day time point, with weaker impacts of rhizobacterial inoculation, suggests that rhizobacteria-induced responses are important to establish as earlier as possible during aphid colonisation. Inoculation of broad bean plants with *Bacillus amyloliquefaciens* altered aphid feeding behaviour, with increased non-probing and salivation events (Serteyn *et al*., 2020), suggesting that aphids may take longer to settle on inoculated plants, and thus longer to initiate reproduction. A combination of behavioural changes and impacts of plant defences on aphid fitness resulting in reduced offspring production are key to reducing aphid populations (Smith & Chuang, 2014). Similarly, any nutritional changes in the plants may lead to imbalances that can impact aphid fitness, but we cannot rule out that the aphid themselves can manipulate the environment to counter plant defensive mechanisms (Züst & Agrawal, 2016). These changes in nutrients might further support activation of signalling pathways and production of defence compounds, as well as boosting the plant growth and development (Sun *et al*., 2020).

We found that both the rhizobacteria *A. radicis* and *B. subtilis* altered the expression of genes in phenylpropanoid biosynthesis pathway. The phenylpropanoid metabolism pathway codes for many different secondary metabolites, and leads to the production of lignin and flavonoids (Vogt, 2010). Phenylpropanoid defensive genes are known to be induced by herbivore feeding and expression varies among resistance and susceptible plant varieties (Ramaroson *et al*., 2022). For example, resistance to aphids in rose plants was linked to biosynthesis of secondary metabolites, including phenylpropanoid, alkaloid and flavonoids (Dong *et al*., 2024). Previous work on *A. radicis* also observed that this rhizobacteria induced the phenylpropanoid and flavonoid pathway in barley linked to microbial N-acyl-homoserine lactone (AHL) production (Han *et al*., 2016). AHLs have been linked to increased plant resilience to stress with correlated changes in root architecture, as well as the plant transcriptome and metabolome (Babenko *et al*., 2022). However, the *B. subtilis* strain does not synthesise AHL molecules (Blake *et al*., 2020), and thus while the shoot expression profiles may be similar between the two rhizobacteria, the mechanisms of effect likely differ. *Bacillus subtilis* interacts with plant roots in various ways including secreting enzymes and phytohormones that trigger systemic resistance in the plant by activating jasmonic acid and salicylic acid signalling pathways (Blake *et al*., 2020). We also observed rhizobacteria-induction of plant phytohormones, including jasmonyl-l-isoleucine synthase encoding for jasmonic acid signalling and several ethylene receptors and transcription factors, confirming these pathways of interest in our system.

A novel result in our study was that rhizobacteria activated arachidonic acid metabolism, which has previously been linked to plant defence to phytopathogens (Dedyukhina *et al*., 2014). We observed very low levels of expression of these genes, and this confirms findings that high arachidonic acid concentration can induce necrosis and accumulation of phytoalexins, while low amounts can elicit systemic and prolonged resistance to phytopathogens (Ivanyuk *et al*., 1990; Ozeretskovskaya, 1994; Dedyukhina *et al*., 2014). Here we speculate that this contributed to our observed microbial-induced resistance against aphids, and thus requires further study

Rhizobacteria inoculation of plant roots can alter plant defences directly, as shown by the various pathways that are up- and down-regulated in inoculated plants compared to controls in our study. Microbial inoculation can also prime plants, where plant defences only activate when triggered by insect feeding (Conrath, 2011; Kim & Felton, 2013). When comparing gene expression of plants inoculated with rhizobacteria without aphids and inoculated plants with aphids, we can begin to disentangle induced and primed responses in the plants. For *A. radicis*, the suppression of phenylpropanoid-related genes occurred independently to aphid feeding, while the induction of glutathione was a primed response activated only on aphid-present plants in the later timepoint when aphid density was high. A study on *Arabidopsis* similarly concluded that the maintenance of glutathione levels played a major role in priming and defending against plant stress (Csiszár *et al*., 2018). For the phenylpropanoid pathway, the increased suppression of genes in this pathway in the later timepoints could be attributed to aphid-induction but the strength of expression varied with rhizobacteria inoculation for both *A. radicis* and *B. subtilis*. This is likely a combination of priming but also a response to the variable aphid densities due to earlier suppression effects. While the population density of herbivores impacts their physiological response, e.g. through crowding effects (Applebaum & Heifetz, 1999). This can also alter their responses to induced plant defences (Frost *et al*., 2008; Züst & Agrawal, 2016). Further experimental work needs to disentangle the effects of insect suppression, that result from microbial-induced plant defences, and the consequences of this reduced aphid density on longer-term colony responses to microbial inoculation.

In conclusion, the rhizobacteria *A. radicis* and *B. subtilis* can suppress aphid populations on barley by inducing and priming a set of plant defensive pathways. These include defence pathways such as phenylpropanoid, glutathione, and phytohormones, as well as several nutritional pathways. Our data suggests that microbial inoculation of barley against aphids is a dynamic phenomenon that acts through various pathways that boost both plant resistance (defences) and tolerance (nutrition and growth) to the growing aphid population. Moreover, we speculate that the strength of these plant defences is modulated by the rhizobacteria in response to aphid density or population expansion. Aphid populations in crop fields tend to be dispersed across multiple host plants, with lower numbers per host unless in event of a local outbreak when number dramatically increase. Therefore, if microbial inoculants can benefit the plant by enabling higher energy allocation to growth for plants when only none or a few aphids are feeding but enable a quick defensive response to increasing aphid densities this can provide efficient crop protection.

## Acknowledgements

Authors acknowledge the project funding by Biotechnology and Biological Science Research Council (BBSRC) of UK [BB/S010556/1]. Further acknowledge the Centre for Genomic Research team at the University of Liverpool, United Kingdom for RNA sequencing work and initial data processing. We also thank Megan Parker and Sophie Blenkinsopp for their support in rearing aphids and maintain the bacterial cultures.

## Data availability

The RNA*seq* data underlying this study will be openly available at the NCBI repository.

Supplementary data may be made available upon request and at the discretion of the authors before peer-review has been completed.

## Author contributions

**SEZ:** conceptualisation of the project idea, obtained grant funding, conducting experiments, experimental data analysis, and results interpretation. **CMM:** experimental design, conducting experiments, processing samples including RNA extraction, transcriptomics data analysis and results interpretation. Both authors wrote the manuscript.

## Supporting Information

**Fig S1.** Gene co-expression network analyses in barley leaves upon rhizobacteria root colonization and shoot herbivory.

**Table S1.** Overview of raw, trimmed and aligned reads.

**Table S2.** Summary of differentially expressed genes in barley leaves upon rhizobacteria root colonization and shoot herbivory.

**Table S3.** Summary of number of genes per enriched KEGG pathways for each treatment and timepoint.

**Table S4.** Enriched KEGG pathways for each treatment group after 24 hours of aphid feeding.

**Table S5.** Enriched KEGG pathways for each treatment group after 7 days of aphid feeding.

**Table S6.** Enriched KEGG pathways for each treatment group after 21 days of aphid feeding.

**Table S7.** Modules of co-expressed genes and number of genes per module in barley leaves upon rhizobacteria root colonization and shoot herbivory.

**Table S8.** Summary statistic of the module trait (treatment group) relationship per timepoint, including after 24 hours, 7 and 21 days of aphid feeding.

**Table S9.** Enriched KEGG pathways for selected module of interest per timepoint, including after 24 hours, 7 and 21 days of aphid feeding.

**Table S10.** The number of hub gene extracted in each selected module of interest per timepoint, after 24 hours, 7 and 21 days of aphid feeding.

**Table S11.** Enriched KEGG pathways for hub genes in the selected module of interest per timepoint, including after 24 hours, 7 and 21 days of aphid feeding.

## References

Applebaum SW, Heifetz Y. 1999. Density-dependent physiological phase in insects. Annual Review of Entomology 44: 317–341.

Babenko LM, Kosakivska I V., Romanenko КО. 2022. Molecular mechanisms of N-acyl homoserine lactone signals perception by plants. Cell Biology International 46: 523–534.

Benjamini Y, Hochberg Y. 1995. Controlling the false discovery rate: A practical and powerful approach to multiple testing. Journal of The Royal Statistical Society. Series B (Methodological) 57: 289–300.

Berendsen RL, Pieterse CMJ, Bakker PAHM. 2012. The rhizosphere microbiome and plant health. Trends in Plant Science 17: 478–486.

Blake C, Christensen MN, Kovács ÁT. 2020. Molecular Aspects of Plant Growth Promotion and Protection by Bacillus subtilis. Molecular Plant-Microbe Interactions® 34: 15–25.

Bray NL, Pimentel H, Melsted P, Pachter L. 2016. Near-optimal probabilistic RNA-seq quantification. Nature Biotechnology 34: 525–527.

Carlson M. 2022.GO.db: A set of annotation maps describing the entire Gene Ontology.

Conrath U. 2009. Priming of Induced Plant Defense Responses. In: Loon LC van, ed. Advances in Botanical Research. Academic Press, 361–395.

Conrath U. 2011. Molecular aspects of defence priming. Trends in Plant Science 16: 524–531.

Csiszár J, Brunner S, Horváth E, Bela K, Ködmön P, Riyazuddin R, Gallé Á, Hurton Á, Papdi C, Szabados L, et al. 2018. Exogenously applied salicylic acid maintains redox homeostasis in salt-stressed Arabidopsis gr1 mutants expressing cytosolic roGFP1. Plant Growth Regulation 86: 181–194.

Dedyukhina EG, Kamzolova S V, Vainshtein MB. 2014. Arachidonic acid as an elicitor of the plant defense response to phytopathogens. Chemical and Biological Technologies in Agriculture 1: 18.

Dong W, Sun L, Jiao B, Zhao P, Ma C, Gao J, Zhou S. 2024. Evaluation of aphid resistance on different rose cultivars and transcriptome analysis in response to aphid infestation. BMC Genomics 25(232): 1–16.

Frost CJ, Mescher MC, Carlson JE, De Moraes CM. 2008. Plant defense priming against herbivores: getting ready for a different battle. Plant physiology 146: 818–824.

Gentleman RC, Carey VJ, Bates DM, Bolstad B, Dettling M, Dudoit S, Ellis B, Gautier L, Ge Y, Gentry J, et al. 2004. Bioconductor: open software development for computational biology and bioinformatics. Genome Biology 5(10): R80.1–R80.16.

Goh CH, Veliz Vallejos DF, Nicotra AB, Mathesius U. 2013. The Impact of Beneficial Plant-Associated Microbes on Plant Phenotypic Plasticity. Journal of Chemical Ecology 39: 826–839.

Han S, Li D, Trost E, Mayer KF, Vlot AC, Heller W, Schmid M, Hartmann A, Rothballer M. 2016. Systemic Responses of Barley to the 3-hydroxy-decanoyl-homoserine Lactone Producing Plant Beneficial Endophyte Acidovorax radicis N35. Frontiers in Plant Science 7 (1868): 1–14.

Huber W, Carey VJ, Gentleman R, Anders S, Carlson M, Carvalho BS, Bravo HC, Davis S, Gatto L, Girke T, et al. 2015. Orchestrating high-throughput genomic analysis with Bioconductor. Nature Methods 12: 115–121.

Ivanyuk VG, Chalova LI, Yurganova LA, Ozeretskovskaya OL, Karavaeva KA. 1990. Immunization of tomatoes by biogenic inductors of protective reactions. Vestn SH Nauki (Moscow*) no* 5: 144–146.

Kim J, Felton GW. 2013. Priming of antiherbivore defensive responses in plants. Insect Science 20: 273–285.

Kolberg L, Raudvere U, Kuzmin I, Vilo J, Peterson H. 2020. gprofiler2-an R package for gene list functional enrichment analysis and namespace conversion toolset g:Profiler.

Langfelder P, Horvath S. 2008. WGCNA: an R package for weighted correlation network analysis. BMC Bioinformatics 9(559): 1–13.

Li Y. 2022. GO/KEGG Enrichment Analysis on Gene Lists from Rice (Oryza Sativa). Bio-protocol 12: 1–12.

Li N, Han X, Feng D, Yuan D, Huang L-J. 2019. Signaling crosstalk between salicylic acid and ethylene/jasmonate in plant defense: Do we understand what they are whispering? International Journal of Molecular Sciences 20(3): 1–15.

van Loon LC, Bakker PAHM, Pieterse CMJ. 1998. Systemic resistance induced by rhizosphere bacteria. Annual Review of Phytopathology 36: 453–483.

Martin M. 2011. Cutadapt removes adapter sequences from high-throughput sequencing reads. EMBnet.journal 17: 10–12.

Martínez-Medina A, Flors V, Heil M, Mauch-Mani B, Pieterse C, Pozo M, Ton J, Dam N, Conrath U. 2016. Recognizing plant defense priming. Trends in Plant Science 21: 818– 822.

Mithöfer A, Boland W. 2012. Plant Defense Against Herbivores: Chemical Aspects. Annual Review of Plant Biology 63: 431–450.

Ozeretskovskaya OL. 1994. Induction of resistance in plants with biogenic elicitors of phytopathogens. Prikladnaia Biokhimiia i Mikrobiologiia 30: 325–339.

Pieterse CMJ, Leon-Reyes A, Van der Ent S, Van Wees SCM. 2009. Networking by small-molecule hormones in plant immunity. Nature Chemical Biology 5: 308–316.

Pineda A, Zheng S-J, van Loon JJA, Dicke M. 2012. Rhizobacteria modify plant–aphid interactions: a case of induced systemic susceptibility. Plant Biology 14: 83–90.

R Core Team. 2022.R: A language and environment for statistical computing.

Ramaroson M-L, Koutouan C, Helesbeux J-J, Le Clerc V, Hamama L, Geoffriau E, Briard M. 2022. Role of Phenylpropanoids and Flavonoids in plant resistance to pests and diseases. Molecules 27(23)8371: 1–24.

Rashid MH-O-, Khan A, Hossain MT, Chung YR. 2018. Induction of Systemic Resistance against Aphids by Endophytic Bacillus velezensis YC7010 via Expressing PHYTOALEXIN DEFICIENT4 in Arabidopsis. Frontiers in Plant Science 9(1904): 1–15.

Ritchie ME, Phipson B, Wu D, Hu Y, Law CW, Shi W, Smyth GK. 2015. limma powers differential expression analyses for RNA-sequencing and microarray studies. Nucleic acids research 43: e47–e47.

Robinson MD, McCarthy DJ, Smyth GK. 2010. edgeR: a Bioconductor package for differential expression analysis of digital gene expression data. Bioinformatics 26: 139–140.

RStudio-Team. 2022. Integrated Development Environment for R. RStudio.

Sanchez-Mahecha O, Klink S, Heinen R, Rothballer M, Zytynska S. 2022. Impaired microbial N -acyl homoserine lactone signalling increases plant resistance to aphids across variable abiotic and biotic environments. Plant, Cell & Environment 45: 3052–3069.

Serteyn L, Quaghebeur C, Ongena M, Cabrera N, Barrera A, Molina-Montenegro MA, Francis F, Ramírez CC. 2020. Induced Systemic Resistance by a Plant Growth-Promoting Rhizobacterium Impacts Development and Feeding Behavior of Aphids. Insects 11; 234: 1–14.

Shikano I, Rosa C, Tan C-W, Felton GW. 2017. Tritrophic Interactions: Microbe-Mediated Plant Effects on Insect Herbivores. Annual Review of Phytopathology 55: 313–331.

Smith CM, Chuang W. 2014. Plant resistance to aphid feeding: behavioral, physiological, genetic and molecular cues regulate aphid host selection and feeding. Pest Management Science 70: 528–540.

Soneson C, Love MI, Robinson MD. 2016. Differential analyses for RNA-seq: transcript-level estimates improve gene-level inferences. F1000Research 4(1521): 1–19.

Sun Y, Wang M, Mur LAJ, Shen Q, Guo S. 2020. Unravelling the roles of Nitrogen nutrition in plant disease defences. International Journal of Molecular Sciences 21:572: 1–20.

Turner TR, James EK, Poole PS. 2013. The plant microbiome. Genome Biology 14(6) 209: 1– 10.

Valenzuela-Soto JH, Estrada-Hernández MG, Ibarra-Laclette E, Délano-Frier JP. 2010. Inoculation of tomato plants (Solanum lycopersicum) with growth-promoting Bacillus subtilis retards whitefly Bemisia tabaci development. Planta 231: 397–410.

Vogt T. 2010. Phenylpropanoid Biosynthesis. Molecular Plant 3: 2–20.

War AR, Paulraj MG, Ahmad T, Buhroo AA, Hussain B, Ignacimuthu S, Sharma HC. 2012. Mechanisms of plant defense against insect herbivores. Plant signaling & behavior 7: 1306–20.

Xi X, Dean A, Zytynska SE. 2024. Microbe-induced plant resistance alters aphid inter-genotypic competition leading to rapid evolution with consequences for plant growth and aphid abundance. Oikos 2024 (6): 1–11.

Züst T, Agrawal AA. 2016. Mechanisms and evolution of plant resistance to aphids. Nature Plants 2: 15206.

Zytynska SE, Eicher M, Rothballer M, Weisser WW. 2020. Microbial-mediated plant growth promotion and pest suppression varies under climate change. Frontiers in Plant Science 11: 1–9.

Zytynska SE, Parker M, Sanchez-Mahecha O. 2024. Rhizobacteria inoculation of plants for reducing insect pests: A meta-analysis on insect behaviour and fitness. bioRxiv: 1–12.

